# Roles of two glutathione *S*-transferases in the final step of the β-aryl ether cleavage pathway in *Sphingobium* sp. strain SYK-6

**DOI:** 10.1101/2020.07.07.192823

**Authors:** Yudai Higuchi, Daisuke Sato, Naofumi Kamimura, Eiji Masai

## Abstract

*Sphingobium* sp. strain SYK-6 is an alphaproteobacterial degrader of lignin-derived aromatic compounds, which can degrade all the stereoisomers of β-aryl ether-type compounds. SYK-6 cells convert four stereoisomers of guaiacylglycerol-β-guaiacyl ether (GGE) into two enantiomers of α-(2-methoxyphenoxy)-β-hydroxypropiovanillone (MPHPV) through GGE α-carbon atom oxidation by stereoselective Cα-dehydrogenases encoded by *ligD, ligL*, and *ligN*. The ether linkages of the resulting MPHPV enantiomers are cleaved by stereoselective glutathione *S*-transferases (GSTs) encoded by *ligF, ligE*, and *ligP*, generating (β*R*β*S*)-α-glutathionyl-β-hydroxypropiovanillone (GS-HPV) and guaiacol. To date, it has been shown that the gene products of *ligG* and SLG_04120 (*ligQ*), both encoding GST, catalyze glutathione removal from (β*R*β*S*)-GS-HPV, forming achiral β-hydroxypropiovanillone. In this study, we characterized the enzyme properties of LigG and LigQ and elucidated their roles in β-aryl ether catabolism. Purified LigG showed an approximately 300-fold higher specific activity for (β*R*)-GS-HPV than that for (β*S*)-GS-HPV, whereas purified LigQ showed an approximately six-fold higher specific activity for (β*S*)-GS-HPV than that for (β*R*)-GS-HPV. Analyses of mutants of *ligG, ligQ*, and both genes revealed that SYK-6 converted (β*R*)-GS-HPV using both LigG and LigQ, whereas only LigQ was involved in converting (β*S*)-GS-HPV. Furthermore, the disruption of both *ligG* and *ligQ* was observed to lead to the loss of the capability of SYK-6 to convert MPHPV. This suggests that GSH removal from GS-HPV catalyzed by LigG and LigQ, is essential for cellular GSH recycling during β-aryl ether catabolism.

**IMPORTANCE:** The β-aryl ether linkage is most abundant in lignin, comprising 45%–62% of all intermonomer linkages in lignin; thus, cleavage of the β-aryl ether linkage together with the subsequent degradation process is considered the essential step in lignin biodegradation. The enzyme genes for β-aryl ether cleavage are useful for decomposing high-molecular-weight lignin, converting lignin-derived aromatic compounds into value-added products, and modifying lignin structures in plants to reduce lignin recalcitrance. In this study, we uncovered the roles of the two glutathione *S*-transferase genes, *ligG* and *ligQ*, in the conversion of GS-HPV isomers, which are generated in the β-aryl ether cleavage pathway in SYK-6. Adding our current results to previous findings allowed us to have a whole picture of the β-aryl ether cleavage system in SYK-6.

## INTRODUCTION

Lignin is one of the major components of plant cell walls, accounting for 15%–40% (1). The industrial production of useful substances from lignin is desired, since it is the most abundant aromatic resource and the second most abundant bioresource on Earth after cellulose. In recent years, microbial conversion processes known as “biological funneling” have attracted attention for their ability to upgrade heterogeneous mixtures of low-molecular-weight aromatic compounds obtained through chemical lignin depolymerization into platform chemicals (2–5). Based on these facts, elucidating the bacterial lignin catabolic system has become more critical.

Lignin is formed through the dehydrogenative polymerization of hydroxycinnamic alcohols, resulting in various C–C and C–O–C bonds between the phenylpropane units (6, 7). Although lignin has several asymmetric carbons, it is considered optically inactive because of its equal amounts of optical isomers (8). Among the intermonomer linkages in lignin, β-aryl ether is the most abundant, comprising 45%–50% of all linkages in softwood lignin and 60%–62% in hardwood lignin (9). Accordingly, cleavage of the β-aryl ether linkage is considered a crucial step in lignin biodegradation. Furthermore, β-aryl ether-type biaryls have two distinct isomeric forms, *erythro* and *threo*, and each form has enantiomeric forms (10, 11).

*Sphingobium* sp. strain SYK-6 is an alphaproteobacterium that exhibits the best-characterized catabolic system for lignin-derived aromatic compounds (3, 12). SYK-6 can utilize lignin-derived biaryls, including β-aryl ether, phenylcoumaran, biphenyl, and diarylpropane, and monoaryls, including ferulate, vanillin, and syringaldehyde, as its sole sources of carbon and energy. In SYK-6 cells, four stereoisomers of a model β-aryl ether compound, guaiacylglycerol-β-guaiacyl ether (GGE), are converted into two enantiomers of α-(2-methoxyphenoxy)-β-hydroxypropiovanillone (MPHPV) through the oxidation of the α-carbon, which is catalyzed by Cα-dehydrogenases, LigD, LigL, and LigN (Fig. 1) (13). LigD oxidizes (α*R*,β*S*)-GGE and (α*R*,β*S*)GGE into (β*S*)-MPHPV and (β*R*)-MPHPV, respectively, whereas LigL/LigN converts (α*R*,β*S*)-GGE and (α*R*,β*S*)-GGE into (β*R*)-MPHPV and (β*S*)-MPHPV, respectively. The ether linkage in the resulting MPHPV is cleaved by enantioselective glutathione *S*-transferases (GSTs), LigF, LigE, and LigP (β-etherases), to produce α-glutathionyl-β-hydroxypropiovanillone (GS-HPV) and guaiacol through the nucleophilic attack of glutathione (GSH) on the MPHPV β-carbon atom (Fig. 1) (14–16). LigF and LigE/LigP attack (β*S*)-MPHPV and (β*S*)-MPHPV, subsequently producing (β*R*)-GS-HPV and (β*S*)-GS-HPV, respectively (14–18). Recently, X-ray crystal structures of LigE and LigF were determined, suggesting that LigE was most similar to the GSTFuA fungal class, and LigF was classified in a new structural class closely related to GSTFuAs or fungal Ure2p-like GST (19). LigF and LigE cleave the β-aryl ether linkage through an *S*_N_2 nucleophilic attack of GSH on the β-carbon of the substrates (18, 19). Another GST, LigG, catalyzes the cleavage of the thioether linkage in (β*R*)-GS-HPV by transferring the (β*R*)-GS-HPV glutathione moiety to another GSH molecule, producing achiral HPV and glutathione disulfide (GSSG) (14, 16). Meux et al. and Pereira et al. independently solved the X-ray crystal structure of LigG (20, 21). LigG belongs to the *omega*-class GST, and Cys15 is a catalytic residue (20). LigG had little to no activity for (β*S*)-GS-HPV, suggesting the involvement of an alternative GST in (β*S*)-GS-HPV conversion (14). A recent study reported that a *Nu*-class GST, NmGST3 of *Novosphingobium* sp. strain MBES04, exhibited GSH-removing activity toward both GS-HPV isomers (22, 23). Kontur et al. reported that NaGST_Nu_, belonging to the *Nu*-class GST, showed activity toward both GS-HPV isomers, and it was the only enzyme essential for converting both GS-HPV isomers in *Novosphingobium aromaticivorans* DSM 12444 (24). Furthermore, they showed that the gene product of SLG_04120, a putative *Nu*-class GST in SYK-6, had GSH-removing activity for both GS-HPV isomers.

**Fig. 1.**
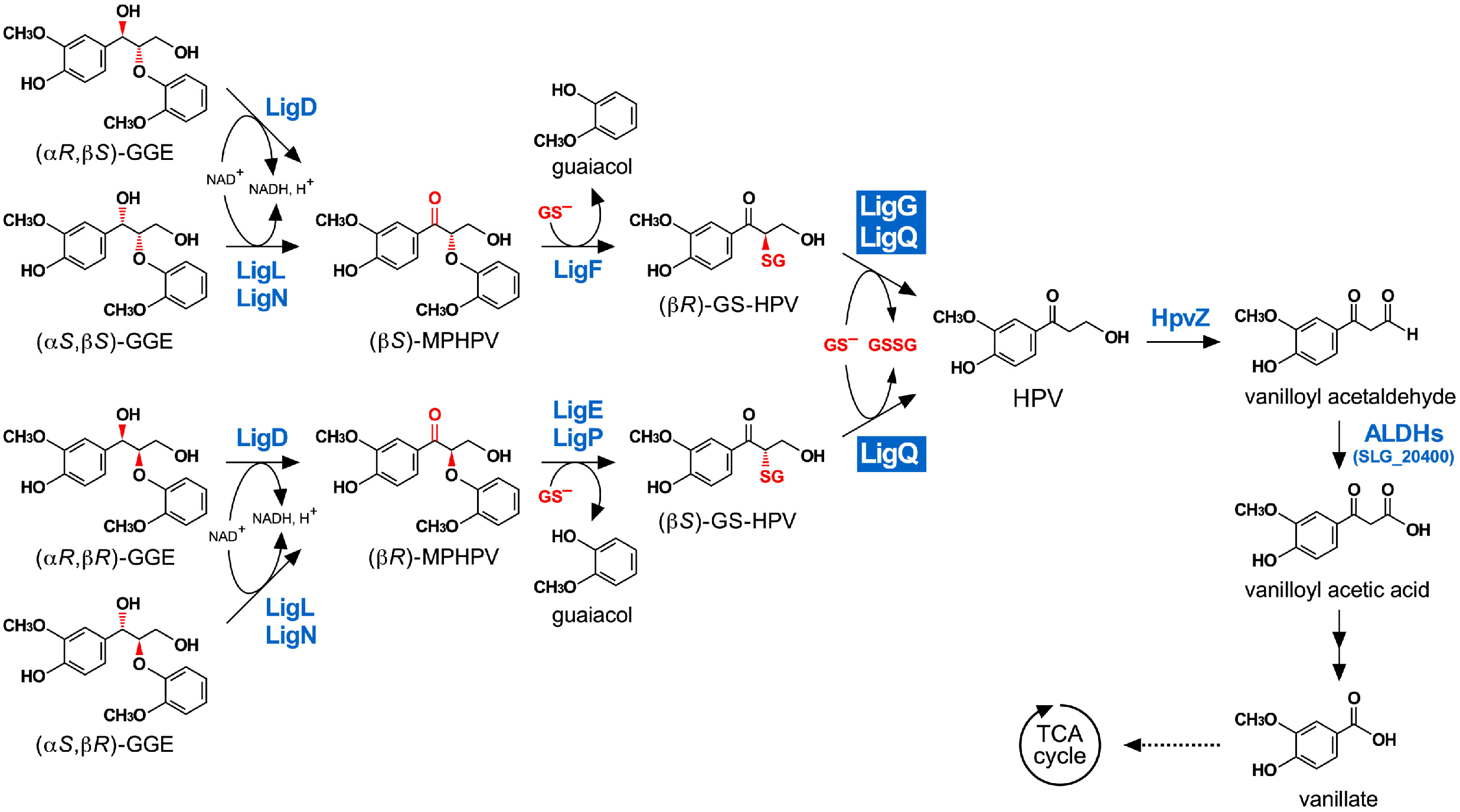
Catabolic pathway of guaiacylglycerol-β-guaiacyl ether in *Sphingobium* sp. strain SYK-6. Stereoisomers of β-aryl ether model compound, GGE, are stereospecifically converted to an achiral C6–C3 monomer, HPV (13–16). The resulting HPV is oxidized to vanilloyl acetic acid via vanilloyl acetaldehyde and further catabolized through vanillate (27, 33). Enzymes: LigD, LigL, and LigN, Cα-dehydrogenases; LigF, LigE, and LigP, β-etherases; LigG and LigQ, GSH-removing enzymes; HpvZ, HPV oxidase; ALDHs, aldehyde dehydrogenases; SLG_20400, vanilloyl acetaldehyde dehydrogenase. *Abbreviations*. GGE, guaiacylglycerol-β-guaiacyl ether; MPHPV, α-(2-methoxyphenoxy)-β-hydroxypropiovanillone; GS-HPV, α-glutathionyl-β-hydroxypropiovanillone; HPV, β-hydroxypropiovanillone; GS^-^, reduced glutathione; GSSG, oxidized glutathione.

In this study, we identified the GS-HPV-converting GST genes in SYK-6 and uncovered the roles of these genes in β-aryl ether catabolism, since the genes involved in GS-HPV catabolism have not been clarified in SYK-6 to date.

## RESULTS

### Preparation of GS-HPV isomers

Racemic MPHPV (400 μM) was reacted with purified LigF and LigE (Fig. S1) with 5 mM reduced glutathione (GSH) to obtain enantiopure GS-HPV isomers. High-performance liquid chromatography (HPLC) analysis of the reaction mixtures showed that MPHPV was converted into GS-HPV by purified LigF and LigE at 60 min (Fig. S2A–C). The conversion of MPHPV by LigF and LigE stopped at approximately 50%, and approximately equimolar GS-HPV to the converted MPHPV was generated (Fig. S2D and E). Chiral HPLC analysis was conducted to confirm whether LigF and LigE enantioselectively convert MPHPV. The MPHPV preparation that was used as the substrate was an equimolar mixture of (β*R*)-MPHPV and (β*S*)-MPHPV (53:47) (Fig. S3A). LigF and LigE completely converted (β*S*)-MPHPV and (β*R*)-MPHPV, respectively, indicating (β*R*)-GS-HPV and (β*S*)-GS-HPV generation in each reaction (Fig. S3B and C). We attempted to remove the remaining MPHPV and guaiacol from the reaction mixtures by ethyl acetate extraction; however, we found that the prepared GS-HPV isomers racemized after incubation in 50 mM Tris-HCl buffer (pH 7.5; buffer A) at 30°C for 24 h (data not shown). Hence, we prepared each GS-HPV isomer by incubating 400 μM MPHPV with purified LigF or LigE and 5 mM GSH before usage.

### Conversion of GS-HPV isomers by *Sphingobium* sp. SYK-6

In order to verify the ability of SYK-6 to convert GS-HPV isomers, the extracts of SYK-6 cells grown in Wx minimal medium (31) containing 10 mM sucrose, 10 mM glutamate, 0.13 mM methionine, and 10 mM proline (Wx-SEMP) were incubated with 100 μM (β*R*)-GS-HPV or (β*S*)-GS-HPV and 2.4 mM GSH. HPLC analysis indicated that SYK-6 converted both GS-HPV isomers into HPV after incubation for 5 min (Fig. S4). The activities of the extracts of SYK-6 cells grown in a Wx-SEMP with or without 5 mM GGE supplementation were measured to evaluate the inducibility of GS-HPV conversion. The specific activity of the extracts from the cells grown in Wx-SEMP for (β*S*)-GS-HPV (142 ± 3 nmol·min^-1^·mg^-1^) was approximately six-fold higher than that for (β*S*)-GS-HPV (24.5 ± 0.2 nmol·min^-1^·mg^-1^). The specific activities for (β*S*)-GS-HPV (150 ± 19 nmol·min^-1^·mg^-1^) and (β*S*)-GS-HPV (31.4 ± 1.1 nmol·min^-1^·mg^-1^) of the extracts from the cells grown in Wx-SEMP supplemented with 5 mM GGE were not significantly higher than those of the extracts from the cells grown in Wx-SEMP. These results suggest that GSH-removing enzyme genes are constitutively expressed.

### Identification of the GST genes involved in GS-HPV isomer conversion

Figure 2 shows a phylogenetic tree of 18 putative GSTs from SYK-6, including LigF, LigE, LigP, LigG, and SLG_04120, with GSTs reported as related enzymes for β-aryl ether catabolism from *Novosphingobium* sp. strain MBES04, *N. aromaticivorans* DSM 12444, *Novosphingobium* sp. PP1Y, and *Thiobacillus denitrificans* ATCC 25259. This phylogenetic tree showed that SLG_00360 is relatively close to a clade, including *ligG* (19% identity), and SLG_04120 belongs to a clade, including NmGST3 of MBES04 (29% identity) and NaGST_Nu_ of DSM 12444 (58% identity), which can convert both GS-HPV isomers. Moreover, Cys15 and Pro16, shown to be the active center of SYK-6 LigG, were conserved in SLG_29340 (Cys10 and Pro11) (Fig. S5). On the basis of these findings, we selected *ligG*, SLG_00360, SLG_04120, and SLG_29340 as candidates for GS-HPV-converting GST genes. To examine whether the gene products of *ligG*, SLG_00360, SLG_04120, and SLG_29340 exhibit activities toward GS-HPV, these genes fused with a His-tag at the 5’ terminus were expressed in *E. coli*. Sodium dodecyl sulfate-polyacrylamide gel electrophoresis (SDS-PAGE) showed the protein production of 33, 28, 36, and 33 kDa, appearing as products of *ligG*, SLG_00360, SLG_04120, and SLG_29340 (Fig. S6). The cell extracts of these *E. coli* transformants were incubated with 100 μM (β*R*)-GS-HPV or (β*S*)-GS-HPV and 2.4 mM GSH. HPLC analysis showed that *ligG* gene product completely converted (β*R*)-GS-HPV into HPV after incubation for 30 min, whereas it only slightly converted (β*S*)-GS-HPV (Fig. 3A and B). This result confirmed that LigG is highly specific for (β*R*)-GS-HPV, which is similar to that of previous reports (14, 16, 21, 25). By contrast, the SLG_04120 gene product completely converted both GS-HPV isomers into HPV, verifying its specificity, as previously reported (Fig. 3C and D) (24). However, SLG_00360 and SLG_29340 gene products showed no activity for both GS-HPV isomers (Fig. 3E–H). Given these results, we designated SLG_04120 as *ligQ* and investigated the role of *ligG* and *ligQ* in converting GS-HPV isomers in SYK-6, as well as the enzymatic properties of these gene products.

**Fig. 2.**
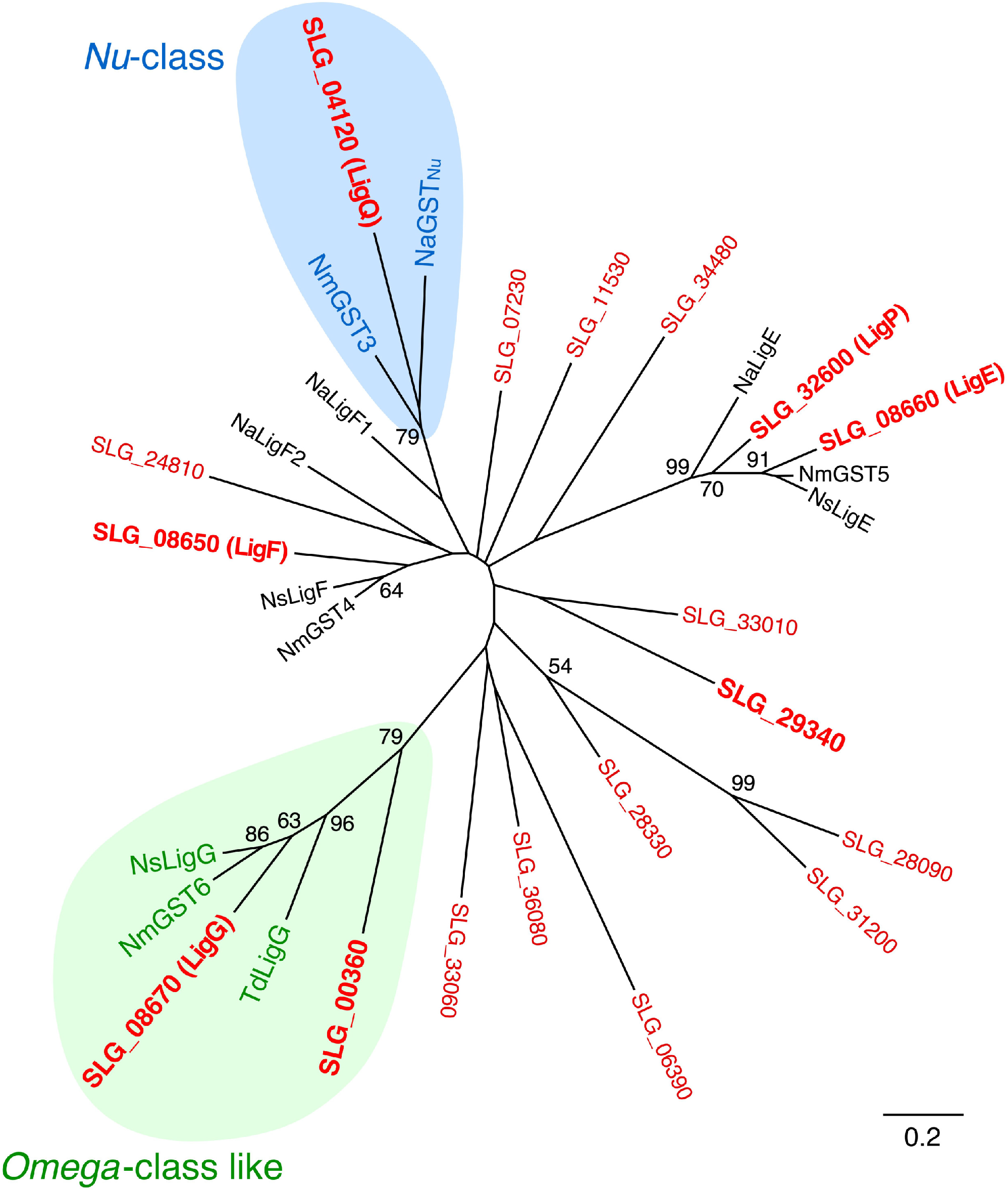
Phylogenetic tree of GSTs from SYK-6 and other bacterial GSTs involved in β-aryl ether conversion. The phylogenetic tree was constructed by neighbor-joining with 1,000 bootstrap replicates. Bootstrap values are indicated at the nodes, and the scale corresponds to 0.2 amino acid substitutions per position. GSTs from SYK-6 are shown in red. *Nu*-class GSTs and *Omega*-class-like GSTs are shown with blue and green backgrounds, respectively. GSTs: LigF (SLG_08650), LigE (SLG_08660), and LigP (SLG_32600), β-etherases of *Sphingobium* sp. SYK-6; LigQ (SLG_04120) and LigG (SLG_08670), GSH-removing enzymes of SYK-6; SLG_00360, SLG_06390, SLG_07230, SLG_11530, SLG_24810, SLG_28090, SLG_28330, SLG_29340, SLG_31200, SLG_33010, SLG_33060, SLG_34480, and SLG_36080, putative GSTs from SYK-6; NmGST4 (GAM05530) and NmGST5 (GAM05531), β-etherases of *Novosphingobium* sp. strain MBES04; NmGST3 (GAM05529) and NmGST6 (GAM05532), GSH-removing enzymes of MBES04; NsLigF (CCA92087) and NsLigE (CCA92088), β-etherases of *Novosphingobium* sp. strain PP1Y; NsLigG (CCA92089), GSH-removing enzyme of PP1Y; NaLigE (ABD26841), NaLigF1 (ABD26530), and NaLigF2 (ABD27301), β-etherases of *Novosphingobium aromaticivorans* DSM 12444; NaGST_N_u (ABD27031), GSH-removing enzyme of DSM 12444; TdLigG (AAZ97003), GSH-removing enzyme of *Thiobacillus denitrificans* ATCC 25259.

**Fig. 3.**
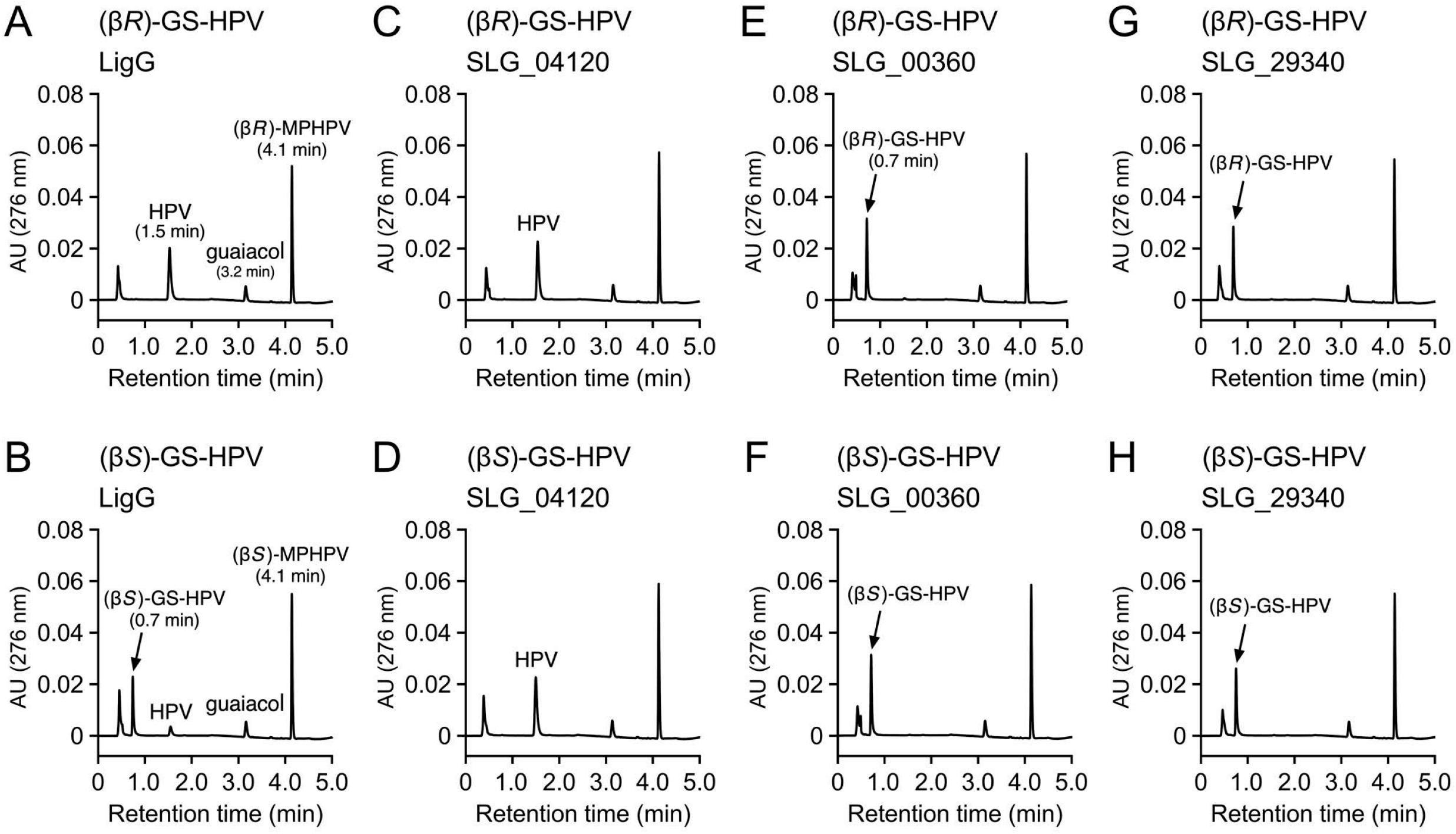
GS-HPV enantiomer conversion by cell extracts of *E. coli* carrying GST genes of SYK-6. (β*R*)-GS-HPV (A, C, E, and G; 100 μM) and (β*S*)-GS-HPV (B, D, F, and H; 100 μM) were incubated with cell extracts of *E. coli* (100 μg protein/ml) harboring each of pET08670 (A and B), pET04120 (C and D), pET00360 (E and F), and pET29340 (G and H) in the presence of 2.4 mM GSH, respectively. Reaction mixtures were collected after incubation for 30 min and analyzed through HPLC.

### Roles of *ligG* and *ligQ* in GS-HPV isomer catabolism

We then created *ligG* mutant (Δ*ligG*), *ligQ* mutant (Δ*ligQ*), and *ligG ligQ* double mutant (Δ*ligG ligQ}* (Fig. S7) to determine whether *ligG* and *ligQ* are indeed involved in converting GS-HPV in SYK-6. The cell extract of each mutant was reacted with 100 μM (β*R*)-GS-HPV or (β*S*)-GS-HPV and 2.4 mM GSH, and their ability to convert GS-HPV isomers was compared with SYK-6. Δ*ligG* and Δ*ligQ* both showed a decreased conversion of (β*R*)-GS-HPV, but only Δ*ligQ* exhibited the loss of (β*S*)-GS-HPV conversion (Fig. 4A and B). Furthermore, Δ*ligG ligQ* had little ability to convert both GS-HPV isomers (Fig. 4A and B). We also examined the activities of Δ*ligG ligQ* cells harboring each of pQF carrying *ligG* (pQF*ligG*) and *ligQ* (pQF*ligQ*) to confirm whether this activity defect was due to the disruption of *ligG* and *ligQ*. A cell extract of Δ*ligG ligQ* harboring pQF*ligG* showed higher specific activity for (β*R*)-GS-HPV than that of the wild type (Fig. 4C). Also, a cell extract of Δ*ligG ligQ* harboring pQF*ligQ* presented almost the same specific activity for (β*R*)-GS-HPV as the wild type (Fig. 4C) and higher specific activity for (β*S*)-GS-HPV than the wild type (Fig. 4D). These results indicated that both *ligG* and *ligQ* participate in the conversion of (β*R*)-GS-HPV in SYK-6 cells, whereas only *ligQ* is involved in (β*S*)-GS-HPV conversion.

**Fig. 4.**
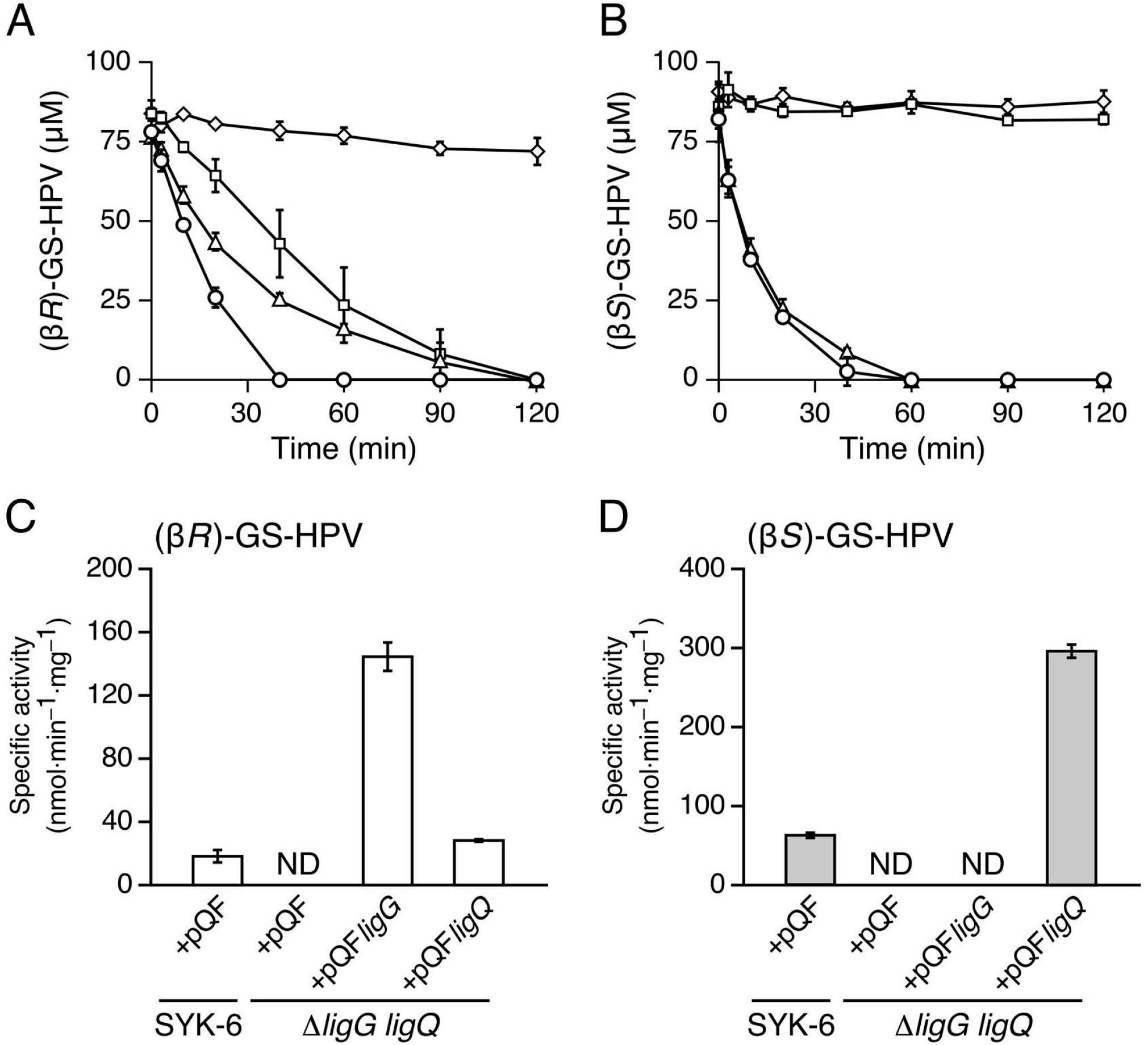
GS-HPV enantiomer conversion by Δ*ligG*, Δ*ligQ*, and Δ*ligG ligQ*. (A and B) Conversion of 100 μM (β*R*)-GS-HPV (A) and (β*S*)-GS-HPV (B) by cell extracts (100 μg protein/ml) of SYK-6 (circles), Δ*ligG* (triangles), Δ*ligQ* (squares), and Δ*ligG ligQ* (diamonds) in the presence of 2.4 mM GSH. (C and D) Complementation of Δ*ligG ligQ* using *pQFligG* and *pQFligQ*, respectively. The cell extracts of SYK-6(pQF), Δ*ligG ligQ*(pQF), Δ*ligG ligQ*(pQF*ligG*), and Δ*ligG ligQ*(pQF*ligQ*) were incubated with 100 μM (β*R*)-GS-HPV (C) and (β*S*)-GS-HPV (D) in the presence of 2.4 mM GSH. All experiments were conducted in triplicate, and each value represents the mean ± standard deviation.

### Enzyme properties of LigG and LigQ

LigG and LigQ were purified to near-homogeneity through nickel affinity chromatography from the cell extracts of *E. coli* expressing His-tagged *ligG* and *ligQ*, respectively (Fig. S8). Purified LigG and LigQ were incubated with 100 μM (β*R*)-GS-HPV or (β*S*)-GS-HPV and 2.4 mM GSH. LigG showed approximately 300-fold higher specific activity for (β*R*)-GS-HPV (33 ± 1 μmolomin^-1^omg^-1^) than for (β*S*)-GS-HPV (0.11 ± 0.01 μmol·min^-1^·mg^-1^). By contrast, LigQ showed approximately six-fold higher specific activity for (β*S*)-GS-HPV (110 ± 14 μmol·min^-1^·mg^-1^) than for (β*R*)-GS-HPV (17 ± 2 μmol·min^-1^·mg^-1^). The specific activity of LigG for (β*R*)-GS-HPV was approximately two-fold higher than that of LigQ. These results indicated that LigG is highly specific for (β*R*)-GS-HPV, and LigQ prefers (β*S*)-GS-HPV more than (β*R*)-GS-HPV. The *K_m_* values of LigG and LigQ for GSH were determined to be 1.19 and 0.34 mM, respectively (Fig. S9).

### Disruption of both *ligG* and *ligQ* affects the MPHPV catabolism

The resting cells of each mutant were reacted with 200 μM racemic MPHPV to investigate whether the disruption of *ligG* and *ligQ* in SYK-6 affects MPHPV conversion. Here Δ*ligG* and Δ*ligQ* cells converted MPHPV similar to the wild type, whereas Δ*ligG ligQ* cells lost the ability to convert MPHPV (Fig. 5A). By contrast, a cell extract of Δ*ligG ligQ* was able to convert MPHPV in the presence of 5 mM GSH at comparable levels with that of the wild type (Fig. S10). The introduction of pQF*ligG* or pQF*ligQ* into Δ*ligG ligQ* cells restored their MPHPV-converting ability, indicating that the disruption of both *ligG* and *ligQ* led to the loss of the ability to convert MPHPV *in vivo* (Fig. 5B).

**Fig. 5.**
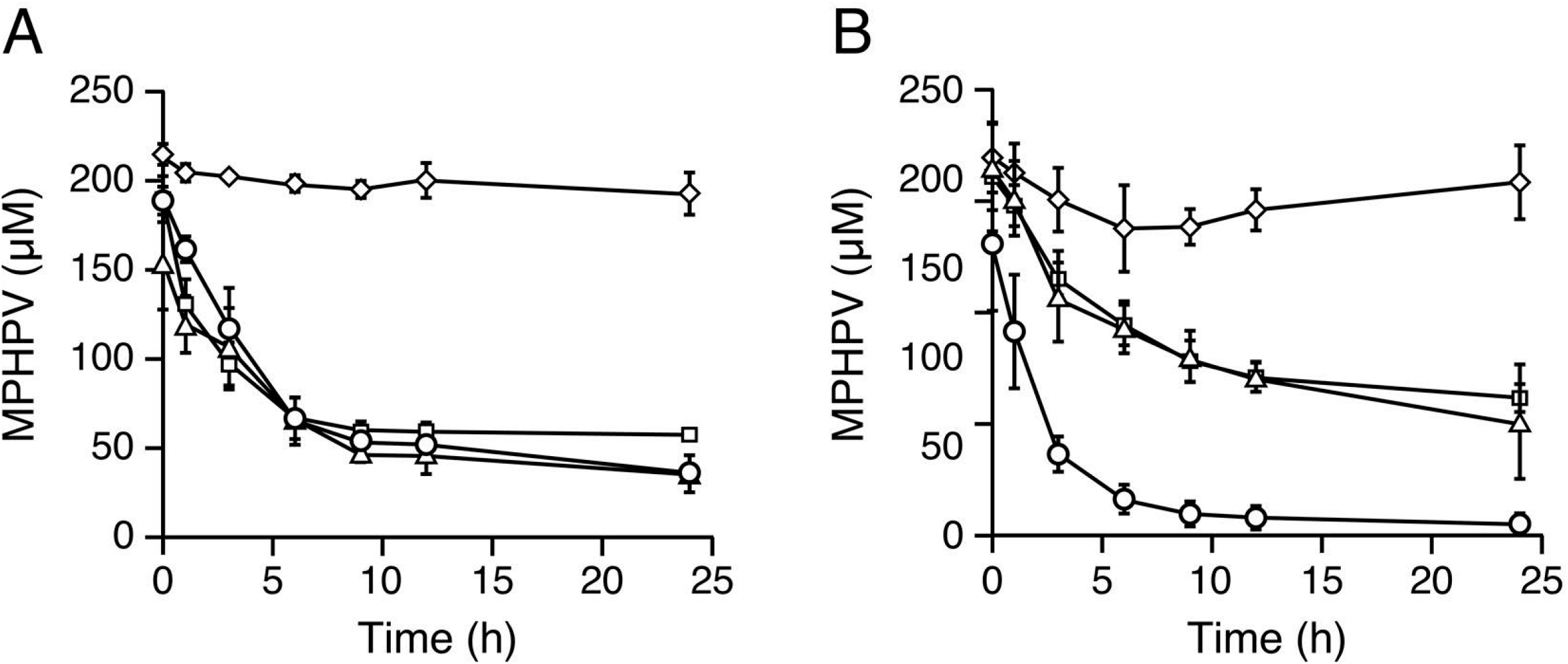
MPHPV conversion by Δ*ligG*, Δ*ligQ*, and Δ*ligG ligQ*. (A) Conversion of 200 μM racemic MPHPV by the resting cells (OD_600_ = 1.0) of SYK-6 (circles), Δ*ligG* (triangles), Δ*ligQ* (squares), and Δ*ligG ligQ* (diamonds). (B) Complementation of Δ*ligG ligQ* using pQF*ligG* and pQF*ligQ*, respectively. Resting cells of SYK-6(pQF) [circles], Δ*ligG ligQ*(pQF) [diamonds], Δ*ligG ligQ*(pQF*ligG*) [triangles], and Δ*ligG ligQ*(pQF*ligQ*) [squares] were incubated with 200 μM racemic MPHPV. All experiments were conducted in triplicate, and each value represents the mean ± standard deviation.

## DISCUSSION

We confirmed that LigG is highly specific for (β*R*)-GS-HPV, and LigQ converts both GS-HPV isomers. Our study further elucidated that both *ligG* and *ligQ* participate in converting (β*R*)-GS-HPV in SYK-6 cells, whereas only *ligQ* is involved in converting (β*S*)-GS-HPV (Fig. 1, 4A, and 4B). Considering that the rate of (β*R*)-GS-HPV conversion by Δ*ligQ* for 60 min was lower than that by Δ*ligG, ligQ* appears to more involved in (β*R*)-GS-HPV conversion than *ligG* (Fig. 4A). However, the specific activity of LigQ for (β*R*)-GS-HPV was approximately two-fold lower than that of LigG in the presence of 2.4 mM GSH. The reason why the disruption of *ligQ* had more influence on (β*R*)-GS-HPV conversion than the disruption of *ligG* may be the higher GSH affinity of LigQ than that of LigG (Table S1, Table S2, and Fig. S9). Alternatively, *ligQ* expression level in SYK-6 may be higher than that of *ligG*.

In *N. aromaticivorans* DSM 12444, NaGST_Nu_, which showed 58% amino acid sequence identity with LigQ, was reported to be the only enzyme essential for the conversion of both GS-HPV isomers. In *Novosphingobium* sp. MBES04, there are NmGST3 and NmGST6 that showed 29% identity with LigQ and 62% identity with LigG, respectively. NmGST3 had activities for both isomers, and NmGST6 was highly specific for (β*R*)-GS-HPV (22, 23); however, the involvement of these genes in GS-HPV catabolism has not been investigated. We found another GST gene in the MBES04 genome, GAM07399, exhibiting 59% and 63% identity with LigQ and NaGST_Nu_, respectively. GAM07399 may be involved in GS-HPV isomer conversion besides the NmGST3 and NmGST6 genes. In *Novosphingobium* sp. PP1Y, NsLigG, exhibiting 63% identity with LigG, was highly specific for (β*R*)-GS-HPV (25). Furthermore, we found CCA92086 in the genome of this strain, which showed 80% identity with NmGST3; and CCA92884 that showed 57% and 66% identity with LigQ and NaGST_Nu_, respectively. These genes may also be involved in GS-HPV isomer conversion. In a phylogenetic tree of LigQ with NaGST_Nu_, NmGST3, GAM07399, CCA92086, CCA92884, known *Nu*-class GSTs (YfcG of *E. coli*, YghU of *E. coli*, and Ure2pB1 of *Phanerochaete chrysosporium*), and putative *Nu*-class GSTs of Sphingomonad, *Nu*-class GSTs were branched into two clades, and LigQ, NaGST_Nu_, GAM07399, and CCA92884 (designated type I) were separated from NmGST3 and CCA92086 (type II) (Fig. S11). Because type I *Nu*-class GSTs are conserved in all strains (SYK-6, DSM 12444, MBES04, and PP1Y), this type of GSTs seemingly plays a major role in GS-HPV conversion.

In the SYK-6 genome, the β-aryl ether catabolic genes, *ligD, ligF, ligE*, and *ligG*, constitute an operon, whereas *ligQ* locates at a different locus (Fig. S12) (14). In the case of MBES04, the NmGST4–NmGST5–NmGST6 genes (GAM05530–05532) corresponding to *ligFEG* are adjacent, and the NmGST3 gene (GAM05529, type II) locates just upstream of GAM05530 with a different transcription direction (Fig. S12) (22). By contrast, the type I *Nu*-class GST gene GAM07399 locates at a different locus. Almost the same gene organization was also observed in the PP1Y genome (Fig. S12) (22). Based on the conservation of these gene arrangements, the conversion of both GS-HPV isomers may become possible by acquiring a *Nu*-class GST gene in strains carrying *ligFEG* or by adding *ligFEG* to a *Nu*-class GST gene-carrying strain. Unlike the aforementioned strains, the genes corresponding to *ligDFE* are scattered through the DSM 12444 genome.

The specific activity of LigG for (β*S*)-GS-HPV (0.11 ± 0.01 μmol·min^-1^·mg^-1^) was only 0.33% of that for (β*R*)-GS-HPV (33 ± 1 μmol·min^-1^·mg^-1^). This result is similar to previous reports demonstrating that LigG had little to no activity for (β*R*)-GS-HPV derivatives (21, 25). Conversely, LigQ showed approximately six-fold higher specific activity for (β*S*)-GS-HPV than for (β*R*)-GS-HPV in this study. Kontur et al. reported that LigQ showed approximately 11-fold higher *k_cat_/K_m_* value for (β*S*)-GS-HPV than for (β*R*)-GS-HPV (Table S2) (24). Also, NaGST_Nu_ of DSM 12444 showed approximately four-fold higher *k_cat_/K_m_* value for (β*S*)-GS-HPV than for (β*R*)-GS-HPV (22), and NmGST3 of MBES04 had approximately two-fold higher specific activity for (β*S*)-GS-HPV than for (β*R*)-GS-HPV (Table S2) (23). Therefore, all these *Nu*-class GSTs have higher specificity for (β*S*)-GS-HPV than for (β*R*)-GS-HPV. The mechanism for the cleavage of the thioether bond in GS-HPV by NaGST_Nu_ was proposed based on X-ray crystal structure, modeling of substrate binding, and mutant analysis. Thr51 and Asn53 of NaGST_Nu_ provide hydrogen bonds that stabilize a reactive glutathione thiolate anion, which attacks the GS^−^ moiety of GS-HPV to form GS–SG disulfide. The rupture of the thioether bond is facilitated by the formation of a transient enolate intermediate. This intermediate is stabilized by the interactions between GS-HPV and Tyr166 and Tyr224 hydroxy groups. The capture of a solvent-derived proton by the carbanion collapses the enolate to form HPV (24). All these residues are conserved in type I *Nu*-class GSTs (LigQ, GAM07399, CCA92884, and YghU), suggesting that these GSTs catalyze glutathione removal from GS-HPV using the same reaction mechanism as that of NaGSTNu.

Finally, we observed that Δ*ligG ligQ* lost the ability to convert MPHPV *in vivo* (Fig. 5A). In Δ*ligG ligQ* cells, intracellular GSH was seemingly depleted due to GS-HPV accumulation during MPHPV catabolism. Consequently, the reactions that are catalyzed by β-etherases (LigF, LigE, and LigP) could not proceed (Fig. 1). Conversion of GS-HPV by LigG and LigQ produces HPV and oxidized glutathione (GSSG); the latter of which must be reduced to GSH by NAD(P)H-dependent glutathione reductase (Fig. 1) (26). Hence, it can be concluded that GS^-^ removal from GS-HPV by LigG and LigQ is essential for maintaining intracellular GSH concentration during β-aryl ether catabolism.

## MATERIALS AND METHODS

### Bacterial strains, plasmids, and culture conditions

Table 1 lists the strains and plasmids that were used in this study. *Sphingobium* sp. strain SYK-6 and its mutants were grown in lysogeny broth (LB), Wx-SEMP, and Wx-SEMP containing 5 mM GGE at 30°C. When necessary, 50 mg kanamycin/liter, 100 mg streptomycin/liter, or 12.5 mg tetracycline/liter were added to the cultures. *E. coli* strains were grown in LB at 37°C. The media for *E. coli* transformants were supplemented with 100 mg ampicillin/liter, 25 mg kanamycin/liter, or 12.5 mg tetracycline/liter.

**Table 1.** Strains and plasmids used in this study.

| Strain or plasmid | Relevant characteristic(s)<sup>a</sup> | Reference or source |
| --- | --- | --- |
| **Strains** |  |  |
| *Sphingobium* sp. |  |  |
| SYK-6 | Wild type; Nal<sup>r</sup> Sm<sup>r</sup> | (34) |
| $\Delta ligQ$ (SME226) | SYK-6 derivative; $\Delta ligQ$; Nal<sup>r</sup> Sm<sup>r</sup> | This study |
| $\Delta ligG$ (SME273) | SYK-6 derivative; $\Delta ligG$; Nal<sup>r</sup> Sm<sup>r</sup> | This study |
| $\Delta ligG ligQ$ (SME274) | SYK-6 derivative; $\Delta ligG ligQ$; Nal<sup>r</sup> Sm<sup>r</sup> | This study |
| *Escherichia coli* |  |  |
| BL21(DE3) | F<sup>-</sup> *ompT hsdS<sub>B</sub>(r<sub>B</sub><sup>-</sup> m<sub>B</sub><sup>-</sup>) gal dcm* (DE3); T7 RNA polymerase gene under the control of the *lacUV5* promoter | (35) |
| HB101 | *recA13 supE44 hsd20 ara-14 proA2 lacY1 galK2 rpsL20 xyl-5 mtl-1* | (36) |
| NEB 10-beta | $\Delta(ara-leu)$ 7697 *araD139 fhuA $\Delta lacX74 galK16 galE15 e14- \phi 80 \Delta lacZ \Delta M15 recA1 relA1 endA1 nupG rpsL$ (Sm<sup>r</sup>) *rph spoT1 $\Delta(mrr-hsdRMS-mcrBC)$** | New England Biolabs |
| **Plasmids** |  |  |
| pRK2013 | Broad-host-range cosmid vector; Km<sup>r</sup> Tc<sup>r</sup> | (37) |
| pET-16b | Expression vector; T7 promoter, Ap<sup>r</sup> | Novagen |
| pAK405 | Plasmid for allelic exchange and markerless gene deletions in *Sphingomonads*; Km<sup>r</sup> | (32) |
| pQF | Expression vector; P<sub>Q5</sub> high-GC-content codon-optimized *cymR*, Tc<sup>r</sup> | (38) |
| pET00360 | pET-16b with a 0.7-kb NdeI-BamHI PCR amplified fragment carrying SLG_00360 | This study |
| pET04120 | pET-16b with a 0.9-kb NdeI-BamHI PCR amplified fragment carrying *ligQ* | This study |
| pET08650 | pET-16b with a 0.8-kb NdeI-BamHI PCR amplified fragment carrying *ligF* | This study |
| pET08660 | pET-16b with a 0.9-kb NdeI-BamHI PCR amplified fragment carrying *ligE* | This study |
| pET08670 | pET-16b with a 0.8-kb NdeI-BamHI PCR amplified fragment carrying *ligG* | This study |
| pET29340 | pET-16b with a 0.7-kb NdeI-BamHI PCR amplified fragment carrying SLG_29340 | This study |
| pAK04120 | pAK405 with a 2.8-kb deletion cassette carrying up- and downstream regions of *ligQ* | This study |
| pAK08670 | pAK405 with a 2.1-kb deletion cassette carrying up- and downstream regions of *ligG* | This study |
| pQF*ligQ* | pQF with a 0.9-kb PCR amplified fragment carrying *ligQ* | This study |
| pQF*ligG* | pQF with a 0.8-kb PCR amplified fragment carrying *ligG* | This study |

### Preparation of substrates

Racemic MPHPV and HPV were prepared as described previously (17, 27). In preparing (β*R*)-GS-HPV and (β*S*)-GS-HPV, purified LigF (40 μg protein/ml) and LigE (150 μg protein/ml) were incubated in 1 ml reaction mixtures containing 400 μM racemic MPHPV, 5 mM GSH, and buffer A, respectively, for 60 min at 30°C. The resulting mixtures were used as preparations of 200 μM GS-HPV isomers, including 4.8 mM GSH. GGE and guaiacol were purchased from Tokyo Chemical Industry, Co., Ltd.

### HPLC analysis

HPLC analysis was conducted using the ACQUITY UPLC system (Waters). The MPHPV and GS-HPV reaction products were analyzed using a TSKgel ODS-140HTP column (2.1 × 100 mm; Tosoh). Additionally, a CHIRALPAK IE column (4.6 × 250 mm; DAICEL) was used to analyze MPHPV reaction products. All analyses were conducted at a flow rate of 0.5 ml/min. The mobile phase was a mixture of solution A (acetonitrile containing 0.1% formic acid) and B (water containing 0.1% formic acid) under the following conditions: detection of MPHPV reaction products using TSKgel ODS-140HTP: 0–2.0 min, linear gradient of 10% A; 2.0–4.0 min, linear gradient from 10% to 60% A; 4.0–4.5 min, linear gradient from 60% to 90% A; 4.5–5.0 min, 90% A. Detection of MPHPV reaction products using CHIRALPAK IE: 0–25.0 min, 30% A. Detection of GS-HPV reaction products using TSKgel ODS-140HTP: 0–5.0 min, 10% A. MPHPV, GS-HPV, guaiacol, and HPV were detected at 280, 286, 276, and 276 nm, respectively.

### Enzyme assays for cell extracts of SYK-6 and its mutants

GS-HPV-transforming activities of the cell extracts of SYK-6 and its mutants were determined by measuring HPV production using HPLC. The MPHPV-transforming activities of the cell extracts of SYK-6 and its mutants were determined by measuring the decrease in MPHPV using HPLC. The reaction products were also detected using HPLC analysis. SYK-6 and its mutant cells (Δ*ligG*, Δ*ligQ*, and Δ*ligG ligQ*) grown in LB were washed with Wx medium, resuspended in Wx-SEMP medium to an optical density at 600 nm (OD_600_) of 0.2, and incubated for 16 h. The resultant cells were washed twice with buffer A. The cells that were resuspended in the same buffer were then broken by an ultrasonic disintegrator (QSonica Q125; WakenBtech Co., Ltd.), and the supernatants were obtained as cell extracts after centrifugation (19,000 ×*g* for 15 min at 4°C). Protein concentration was determined using the Bradford method, using bovine serum albumin as the standard (BioRad Laboratories).

Cell extracts (5–1,000 μg protein/ml) were incubated with 100 μM (β*R*)-GS-HPV, (β*S*)-GS-HPV, or 200 μM racemic MPHPV in the presence of 2.4 mM (for GS-HPV) or 5.0 mM GSH (for MPHPV) in buffer A at 30°C. The reactions were stopped by adding acetonitrile (final concentration of 50%) after 3 min (measurement of the specific activities for GS-HPV), 5 min (identification of GS-HPV metabolites), 20 min (measurement of the specific activities for GS-HPV in mutant-complemented strains), or 60 min (for MPHPV). Precipitated proteins were removed by centrifugation at 19,000 ×*g* for 15 min, and the resulting supernatants were diluted with water to a final acetonitrile concentration of 12.5%, filtered, and analyzed using HPLC. The specific activity of GS-HPV conversion was expressed in moles of HPV produced per min per milligram of protein. The cells of SYK-6 grown in LB were washed with Wx medium, resuspended in Wx-SEMP medium to an OD_600_ of 0.2, and incubated at 30°C to examine the effects of GGE on enzyme induction. When the cultures reached an OD_600_ of 0.4–0.5, we added 5 mM GGE to them. After 12 h of further incubation, cell extracts were prepared and used for the enzyme assay. For complementing Δ*ligG ligQ*, pQF*ligG* and pQF*ligQ* were constructed by In-Fusion cloning of *ligG* and *ligQ* fragments amplified by PCR using SYK-6 total DNA and their corresponding primer pairs (Table 2) into pQF. After confirming the nucleotide sequences of the inserts, each plasmid was introduced into Δ*ligG ligQ* cells through electroporation. The transformed cells were grown in Wx-SEMP containing 100 μM cumate and tetracycline for 16 h, and the cell extracts were prepared and used for the enzyme assay.

**Table 2.**
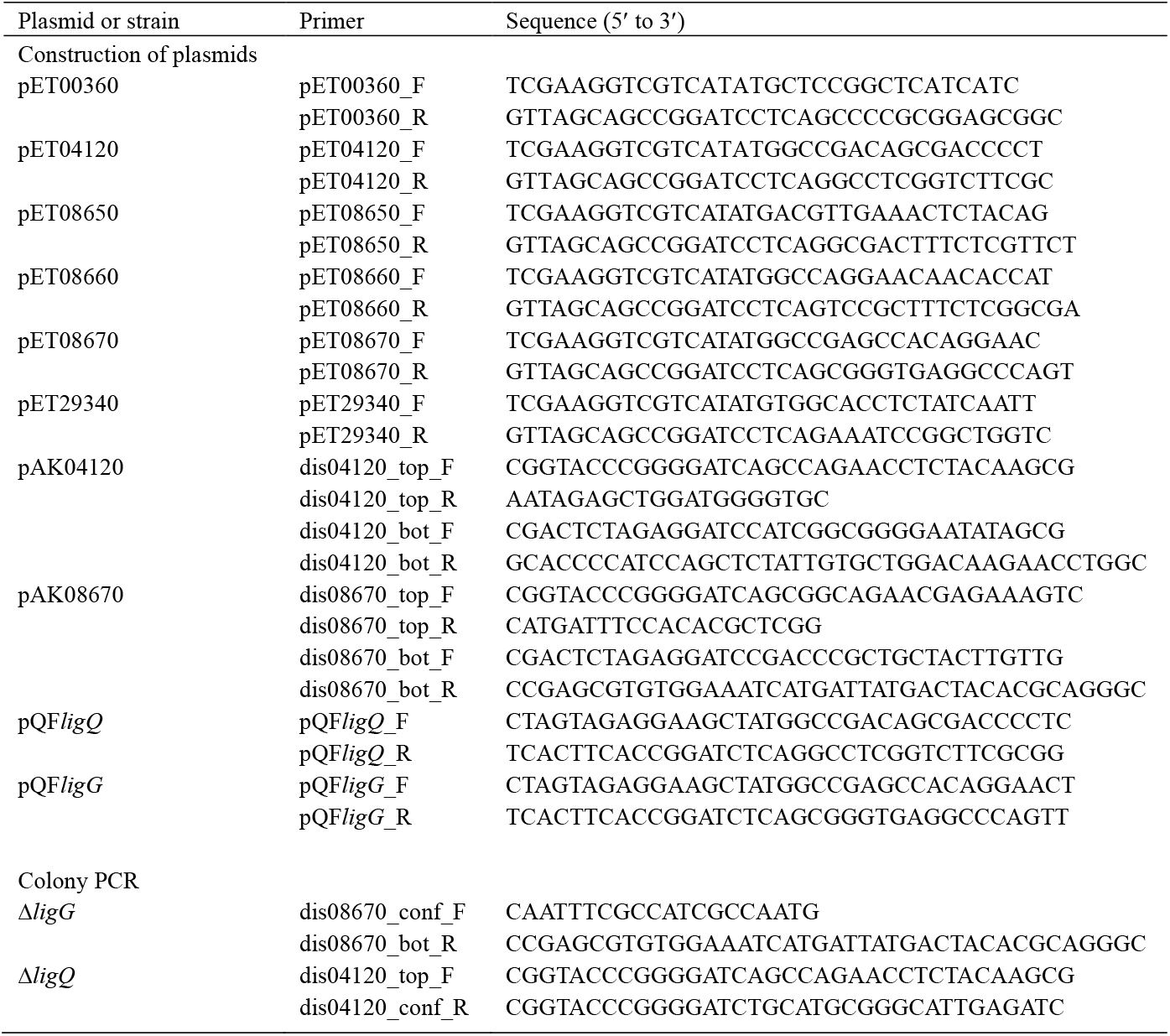
Primers used in this study.

### Sequence analysis

Nucleotide sequences were determined through Eurofins Genomics. Sequence analysis was conducted using the MacVector program (MacVector, Inc.). Sequence similarity searches, pairwise alignments, and multiple alignments were conducted using the BLASTP program (28), the EMBOSS Needle program through the EMBL-EBI server (29), and the Clustal omega program (30), respectively. For phylogenetic analysis, multiple alignments were conducted using the Clustal omega program, and then, phylogenetic trees were generated using the neighbor-joining algorithm of the MEGA 7 software (31), applying 1,000 bootstrap replicates.

### Expression of SYK-6 GST genes in *E. coli* and enzyme purification

DNA fragments carrying SLG_00360, *ligQ* (SLG_04120), *ligF* (SLG_08650), *ligE* (SLG_08660), *ligG* (SLG_08670), and SLG_29340 were amplified through PCR using SYK-6 total DNA and the primer pairs listed in Table 2, and each amplified fragment was cloned into pET-16b by In-Fusion Cloning (Takara Bio). The nucleotide sequences of the inserts were confirmed by sequencing. Each expression plasmid was introduced into *E. coli* BL21(DE3), and the transformed cells were grown in LB. Gene expression was induced for 4 h at 30°C by adding 1 mM isopropyl-β-D-thiogalactopyranoside when the OD_600_ of the cultures reached 0.5. The cells were then harvested by centrifugation at 5,000 *×g* for 5 min at 4°C and washed with buffer A. The cells were then resuspended in the same buffer and broken using an ultrasonic disintegrator. The supernatants were obtained as cell extracts after centrifugation at 19,000 ×*g* for 15 min at 4°C. For purification of LigF, LigE, LigG, and LigQ, the cell extracts of *E. coli* BL21(DE3) harboring pET08650, pET08660, pET08670, and pET04120, respectively (Table 1), were transferred onto a His SpinTrap column (GE Healthcare). The purified fractions were subjected to desalting and concentration using an Amicon Ultra centrifugal filter unit (30 kDa cutoff; Merck Millipore), and the enzyme preparations were stored at −80°C. Gene expressions and the purity of the preparations were examined using SDS-12% PAGE. The protein bands in gels were stained with Coomassie Brilliant Blue.

### Enzyme assays for cell extracts of *E. coli* transformants and purified enzymes

GS-HPV-transforming activities were determined by measuring HPV production using HPLC. The reaction products were also detected through HPLC analysis. The cell extracts (100 μg protein/ml) and purified enzymes (LigG, 1.25–250 μg protein/ml; LigQ, 0.2–1.0 μg protein/ml) were incubated in the 100 μl reaction mixtures containing 100 μM (β*R*)-GS-HPV or (β*S*)-GS-HPV, 2.4 mM GSH, and buffer A at 30°C. The reactions were stopped by adding acetonitrile or methanol to a final concentration of 50% after 30 min of incubation (for cell extracts) and 15 s (for purified enzymes). Protein precipitates were removed through centrifugation at 19,000 *×g* for 15 min. The resulting supernatants were diluted with water to a final concentration of acetonitrile or methanol of 12.5%, filtered, and analyzed using HPLC. Specific activities for GS-HPV conversion of purified enzymes were calculated after 15 s incubation and expressed in moles of HPV produced per min per milligram of protein. The *K_m_* values of LigG and LigQ for GSH were determined using 100 μM (β*R*)-GS-HPV under the following protein and GSH concentrations: LigG, 1.25 μg/ml protein and 0.25–4.9 mM GSH; LigQ, 5.0 μg/ml protein and 0.1–1.9 mM GSH. The *K_m_* values were calculated through non-linear regression analysis using the GraphPad Prism 7 software (GraphPad Software Inc.) fitted to the Michaelis–Menten equation and expressed as means ± standard deviations of three independent experiments.

### Construction of mutants

To construct the *ligG* and *ligQ* mutants, the upstream and downstream regions of the genes were amplified through PCR from SYK-6 total DNA using the primer pairs listed in Table 2. The resulting fragments were cloned into pAK405 by In-Fusion Cloning. Each of the resulting plasmids was introduced into the SYK-6 cells by triparental mating, and the resulting mutants were selected as described previously (32). Gene deletion was confirmed through colony PCR using the primer pairs listed in Table 2.

### Resting cell assay

The cells of SYK-6, Δ*ligG*, Δ*ligQ*, and Δ*ligG ligQ* were grown in Wx-SEMP for 16 h. The cells were then collected by centrifugation, washed twice with buffer A, and resuspended in the same buffer. After adding 200 μM of MPHPV, the resting cells (OD_600_ of 1.0) were incubated at 30°C with shaking for 24 h. Portions of the cultures were periodically collected, and the amounts of the MPHPV were measured using HPLC. To complement Δ*ligG ligQ*, the resting cells (OD_600_ of 5.0) of SYK-6 harboring pQF, Δ*ligG ligQ* harboring pQF, Δ*ligG ligQ* harboring pQF*ligG*, or Δ*ligG ligQ* harboring pQF*ligQ*, grown in Wx-SEMP containing cumate (100 μM) and tetracycline, were incubated with 200 μM MPHPV at 30°C with shaking for 24 h.

## ACKNOWLEDGMENT

This work was supported in part by a research grant from the Uchida energy science promotion foundation.

